# Protein-protein interaction is a major source of epistasis in genetic interaction networks

**DOI:** 10.1101/2024.11.11.623081

**Authors:** Xavier Castellanos-Girouard, Adrian WR Serohijos, Stephen W Michnick

**Affiliations:** Department of Biochemistry, Université de Montréal; Montreal, H3T 1J4, Canada; Robert-Cedergren Center for Bioinformatics and Genomics, Université de Montréal; Montreal, H3T 1J4, Canada

## Abstract

Genetic interaction (GI) and protein-protein interaction (PPI) networks have proved among the most important tools for inference of gene function and genotype-phenotype relationship^1–6^. It remains unclear, however, how these two networks are related to each other. Here we demonstrate that the strengths of epistatic genetic interactions between two genes strongly correlate with the binding free energies of interactions between their cognate proteins, for both yeast and human genomes. Consequently, we show that GI and PPI networks can reciprocally predict each other. Further, we observe that functional divergence and redundancy in duplicated genes (paralogs) are reflected in the binding affinity of their interactions with other proteins and in their genetic interaction strengths. Finally, we demonstrate that the overall topologies of GI and PPI networks significantly overlap in two topological features: Modules, in which genes/proteins are organized by common function, and connectors, which link modules to each other. This opens avenues for new integrated network approaches in understanding cellular processes and flow of genomic information.

## Introduction

Determining how genes cooperate to shape phenotype is a central goal in molecular biology. At the cellular level, our understanding of functional relationships between genes is best reflected in genetic interaction (GI) and protein-protein interaction (PPI) networks^1,3^. These two networks are highly complementary; GI networks link broad functional relationships between genes via their contribution to a phenotype, whereas PPI networks are maps of physical contacts between proteins. However, how these two networks are related to each other remains an open question^1,7^.

GI networks are based on systematic or pooled pairwise deletion or disruption of genes and a genetic interaction is scored as effects on some trait, such as cell proliferation rates, that are less than or greater than the product of the effects of deleting the individual genes, a phenomenon called epistasis^8,9^. Epistasis is a major source of complexity in predicting genotype-phenotype relationships, making the elucidation of its molecular mechanism a significant endeavor^2,10–13^. GI networks also serve important applications such as identifying unique genetic vulnerabilities in cancer cells and inferring gene function by organizing genes into functional groups^1,14–16^. PPIs are also useful for inferring gene functions, linking proteins of unknown, to those of known function^3,17^. Finally, about two-thirds of mutations that cause Mendelian human diseases also disrupt PPIs, linking the integrity of PPI networks to disease^18^. Considering their central role in understanding normal and pathological cellular processes, modelling the relationship between GI and PPI networks has been a longstanding goal in systems biology^7,19^.

Recent quantitative proteomic studies provide clues of a deeper physical relationship between PPI and GI networks^20–22^. The number of complexes formed between two proteins is determined by their abundances and their binding affinity. Hein et al. used mass spectrometry to measure the ratios of the abundances of the two binding partners (abundance stoichiometry) and of total numbers of protein pairs that form complexes (interaction stoichiometry), a number proportional to binding dissociation constants (K_d_)^20^. The ranges of these binding and abundance stoichiometries were remarkably broad, between ∼10^−7^-10^1^ (Fig. 1d, regions 2 and 4)^20^. Additionally, well annotated and known stable protein complexes were shown to be enriched around 1:1 abundance and interaction stoichiometries, suggesting a balance of the relative concentration of proteins that form such stable PPIs. The strong negative epistasis that can occur among genes that code for the subunits of essential and stable protein complexes raises the question of whether the broad range of binding stoichiometries could be proportional to epistatic strengths^1,7^.

**Figure 1.**
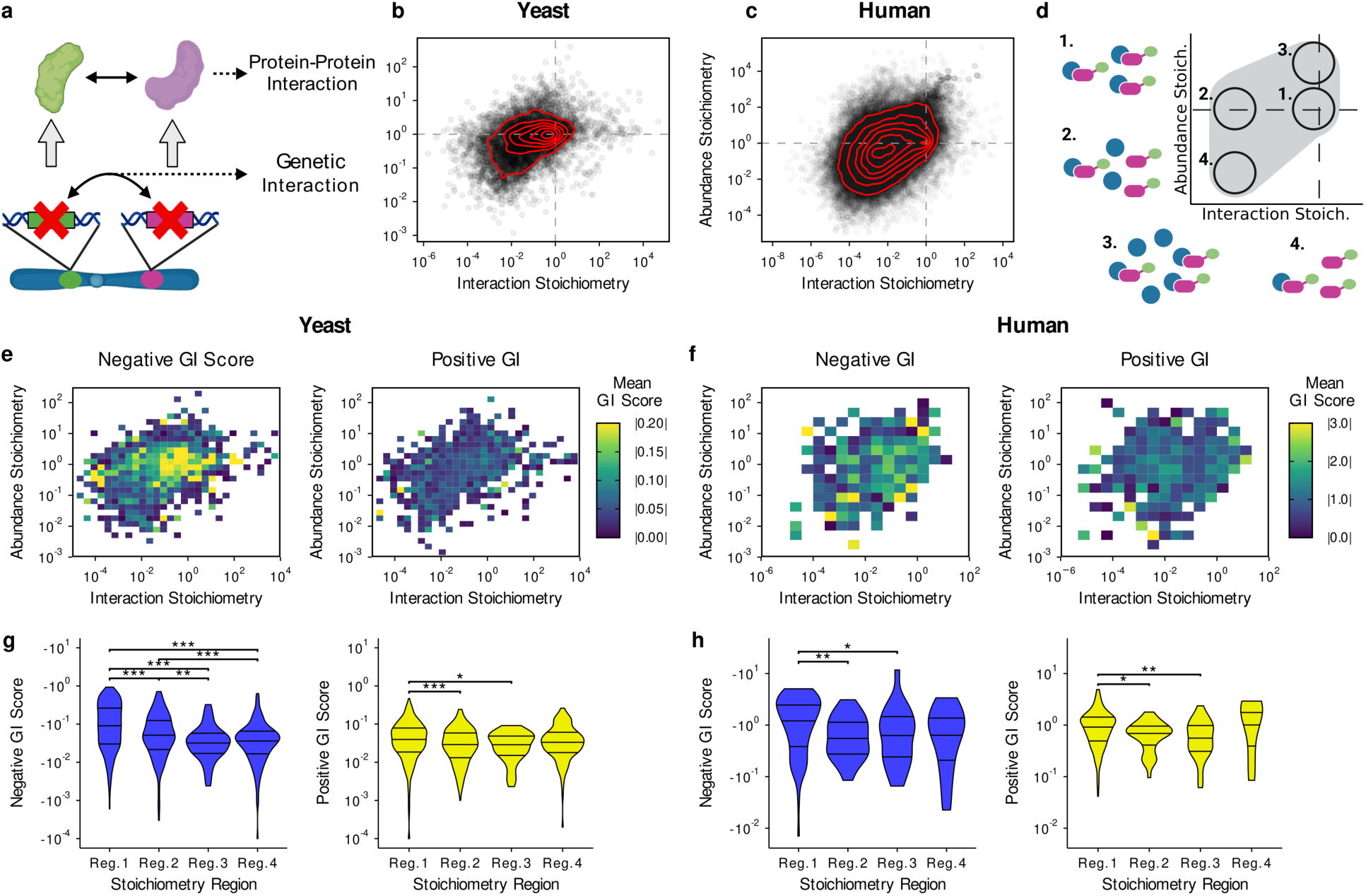
Strong negative GIs map to PPIs of equal protein stoichiometry. **a**, Genetic interactions are generated by comparing fitness defects in genotypes with single or double gene knockout/knockdowns, and protein-protein interactions are defined as physical contacts between proteins. Illustration created with BioRender. **b, c**, Protein-protein interactions (grey points) are plotted according to two stoichiometries: Interaction Stoichiometry and Abundance Stoichiometry for yeast (b) and human cell lines (c). **d**, The stoichiometry plot can be separated into 4 regions representing PPIs with different ranges of protein stoichiometry values. Based on figure from reference 20. Illustration created with BioRender. **e (yeast), f (human),** Mean negative and positive genetic interaction scores for PPIs throughout the stoichiometry plot. GI scores in human were taken from a screen in K562 cell lines. **g, (yeast), h, (human),** Distribution of GI scores within the 4 defined regions of the stoichiometry plot. Significance levels are annotated as: * for *p* ≤ 0.05, ** for *p* ≤ 0.01, and *** for *p* ≤ 0.001.

In addition, Cho et al. demonstrated that a quantitative measure of cellular spatial organization similarity, measured by spinning disk confocal microscopy, of interacting proteins is proportional to interaction stoichiometry^21^. These results imply that higher order physical organization of proteins in the cell is also determined by binding strength. This relationship also defines topological features of cellular PPI networks in which high interaction stoichiometry PPIs formed modules of functionally related proteins while the low interaction stoichiometry pairs formed transient interactions that connect modules^20,21^. GI networks also show a highly modular organization and the correspondence of modular structures between PPI and GI networks has been established^1,7,23^. Is it possible that there is a similar correspondence of PPI protein and GI pairs in connector regions of PPI and GI networks?

Finally, it has been demonstrated that common patterns of epistasis between pairs of mutations in subunits of protein complexes and a set of gene knockout strains correlate with the distance between the mutated residues in protein complexes^24–26^. Consistent with the notion that mutations of residues within functional sites, including hydrophobic core, allosteric, catalytic or binding sites, should produce similar phenotypes and equally, be more strongly energetically coupled to each other^27–29^.

Following from these three observations, here, we test the hypotheses that the strengths of PPI and GI correlate and that PPI networks and GI networks overlap in their topological features (modules and connectors).

## Results

### Epistasis varies with protein abundance and PPI stoichiometries

Since a complete systematic GI network has been constructed for the budding yeast *Saccharomyces cerevisiae* (henceforth “yeast), we chose to make our initial comparison of PPI binding and epistatic strengths for this organism^1^. We first constructed a protein abundance versus PPI binding protein stoichiometry map based on data from a recent affinity purification-mass spectrometry and a meta-analysis of protein abundance measurements^20–22,30^. The distribution of yeast PPIs in the stoichiometry plot (Fig. 1b, Supplementary data 1) is qualitatively similar to that of human proteins (Fig. 1c, Supplementary data 2), the major difference being a higher frequency of low stoichiometry interactions for human proteins. This higher frequency of weak interactions could reflect relaxation of selection pressure that occurred in the transition from unicellular to multicellular organisms as a consequence of reduced population, allowing for the increased rate of emergence of new genomic features, such as gene duplicates and accrual of new gene functions^31–33^. It could also reflect the increase in requirements of connector PPIs to coordinate activities of the expanded number of functional modules in metazoans. Furthermore, we also found that region 1 (Fig. 1b, d) in the yeast stoichiometry plot is enriched for stable complexes (Extended Data Fig. 1), consistent with findings for human proteins^20^. Notwithstanding the evolutionary distance between the two organisms, the similarity in their stoichiometry maps may reflect the conservation of core cellular processes, particularly the molecular mechanisms that mediate them, across yeast and humans^34^.

We next sought to explore how GIs distribute across the PPI stoichiometry map. To this end, we merged the GI and PPI networks using the criterion that if the genes comprising a GI also code for proteins that interact, the GI score was assigned to that PPI (Supplementary Data 2, 3). Yeast GIs were taken from a near-complete digenic knockout screen^1^ and human GIs were taken from the largest double knockdown study in human cell lines (Jurkat and K562) to date^35^. Of note, the measured GI network in human cell lines is only comprised of 472 genes and consequently only about 0.05% of all possible double knockouts were measured, a much smaller sample than the data for yeast. By visualizing average GI strengths throughout the stoichiometry plots, we observed that negative GIs are strongest for PPIs near 1:1 interaction and abundance stoichiometries (Fig. 1e, left panel). This observation is consistent with previous analyses in yeast showing that negative GIs are enriched between genes whose proteins assemble into complexes^1^. Some examples are highlighted in Extended Data Table 1. This pattern also holds in human cells (Fig. 1f, left panel, Extended Data Fig. 2a), but to a lesser extent, likely due to the smaller sample size. In contrast, the strength of positive GIs seems to be distributed more uniformly across the stoichiometry map. This suggests that positive GIs are not predominantly driven by PPI and may be better explained in combination with other mechanisms. Indeed, suppression mechanisms involving mRNA decay and protein degradation, as well as specific patterns of disruption in metabolic pathways have been proposed as important sources of positive GIs^1,36–38^.

To further investigate the distribution of GI strength throughout the stoichiometric landscape, we divided interactions into 4 regions previously outlined by Hein et al. (Fig. 1d): region 1 consists of interacting protein pairs of equal abundances and 1:1 binding ratios (enriched for stable complexes); region 2 consists of protein pairs of equal abundances but low binding ratios (enriched for transient interactions); region 3 consists of protein pairs of unequal abundances and 1:1 binding ratios, and region 4 consists of protein pairs of unequal abundances and low binding ratios^20^. As suggested by the stoichiometry plots, the distribution of negative GIs within region 1 is significantly higher than that of other regions (P < 0.01, Wilcoxon rank-sum test) (Fig. 1g, h, left panels, Extended Data Fig. 2c). The distribution of positive GI values is also highest in region 1 in yeast and human cell lines (Fig. 1g, h, right panels), in agreement with previous observations that complexes, composed mostly of non-essential genes, are enriched for positive GIs (Extended Data Table 2)^1^. Interestingly, region 3 that is characterized by high interaction stoichiometry and an excess of affinity purified versus bait protein show the lowest number of GIs among the four regions (Fig. 1g and h, and Extended Data Fig. 2c), reflecting that the number of protein complexes formed is not the sole determinant of GI strength. Taken together, these results show that there is robust overlap between GI and PPI networks.

### Negative epistasis correlates with PPI binding strength

Substantial efforts have been made to model how the physicochemical properties of proteins such as folding stability and PPI binding energies drive epistasis^29,39,40^. Whilst these approaches explain how epistasis arises from point mutations for single proteins or pairs of proteins in PPIs, the extent to which the strengths of GIs for whole gene knockout (or knock down) could be explained by binding free energies is unknown.

Interaction stoichiometry can provide an indirect measure of PPI strength if protein pairs are of equal abundance, however, interaction stoichiometry is inherently limited by the abundance of the bait in an affinity purification-mass spectrometry (AP-MS) experiment^20,21^. This is reflected in the triangular shape of stoichiometry maps (Fig. 1b to d), where protein pairs of very low abundance stoichiometry have a strong tendency to not form high interaction stoichiometry interactions (Fig. 1b and c). Abundance is therefore a confounding factor when using interaction stoichiometry as a proxy for binding strength.

To obtain direct measures of PPI binding affinities, we devised a simple approach to estimate equilibrium dissociation constants (K_d_) from protein stoichiometries and protein abundances measured by mass spectrometry. Briefly, the number of complexes formed by a pair of proteins was estimated using the measured interaction stoichiometry and cellular abundances. The number of complexes were then used to estimate the number of unbound copies of each interactor. Finally, copy numbers were converted to concentrations, from which second order equilibrium dissociation constants (K_d_) were calculated (Supplementary Data 5, 6). We validated our results by comparing the estimated values to *in vitro* measurements of dissociation constants from literature and found a moderate but significant correlation of 0.40 (Pearson’s r, *p*-value = 5.2 × 10^−4^, Extended Data Figure 3a, Supplementary Data 7), which was comparatively better than that of interaction stoichiometry (Pearson’s r = −0.29, *p*-value = 0.016, Extended Data Fig. 3b). The theoretically expected relationship between K_d_ and interaction stoichiometry is inversely proportional. It has been noted that important criteria to assess reliability (*i.e.*, controlling for titration and demonstrating equilibrium) in equilibrium binding experiments often go unreported throughout literature, and different experimental approaches can sometimes yield non-overlapping values of K_d_^41,42^. Moreover, experiments are generally performed *in vitro* and under conditions that do not reflect native cellular environments. Considering these important sources of variability, we reasoned that our values fall within an acceptable range of precision. To our knowledge, this is the first report of a proteome-wide estimate of *in vivo* binding affinities.

A density heat map relating GI scores and K_d_ shows a visible bias towards negative GIs for high affinity interactions in yeast and human (Extended Data Fig. 4a). To quantitatively evaluate the relationship between K_d_ and GI strength, we binned the PPIs into quantiles according to K_d_ and calculated the mean GI strength within each bin (Extended Data Fig. 4b, c). The mean strengths of negative GIs correlate remarkably and significantly with K_d_ for both yeast (Fig. 2a, Spearman’s ρ = −0.98, *p*-value = 2.2 × 10^−5^) and human PPIs (Fig. 2b, Spearman’s ρ = 0.95, *p*-value = 0.014). A weak but significant correlation is also observable for positive GIs in yeast (Fig. 2c, Spearman’s ρ = 0.76, *p*-value = 0.008), which is likely driven by an enrichment of significant positive GIs within complexes composed of non-essential genes (Extended Data Fig. 5c, Extended Data Table 2)^1,7^. At the molecular level, this trend could be explained by a mechanism where the absence of a single protein (single gene knockout) is enough to completely disrupt the assembly of a complex in which it is embedded^43^. In this case, the growth defect for the double knockout would be expected to be less than that of the product of the two single knockout. Specifically, in protein biochemistry, the presence of a binding partner can increase the folding stability of a protein by providing nucleation sites for its unfolded species, an effect that is heightened for stronger PPIs^44^. The single gene knockout could destabilize the other gene, consequently, the double knockout is not substantially more drastic than the single knockout.

**Figure 2.**
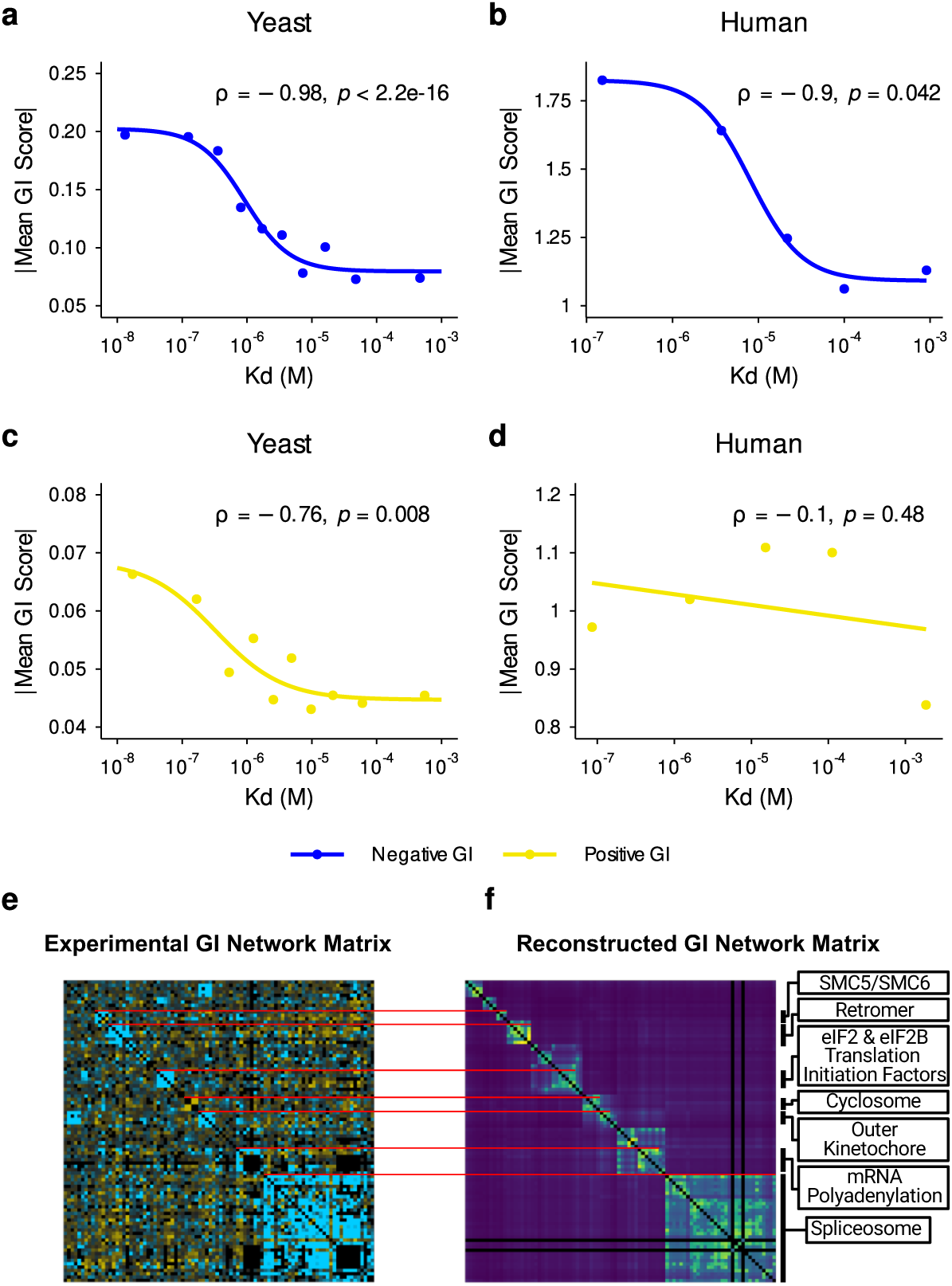
Genetic interactions correlate with PPI binding affinity. **a**, Binning PPIs according to K_d_s reveals a significant monotonic relationship between K_d_ and negative GIs. PPIs in yeast were binned into deciles (10 quantiles). **b**, The binning analysis described in (a) was repeated for negative genetic interaction data from K562 human cell lines. Data were binned into quintiles (5 quantiles) to adjust for data sparsity. **c**, **d**, The binning analyses described in (a) and (b) were performed on positive genetic interactions for yeast (c) and the human K562 cell line (d). The GI versus K_d_ trend was fit to a sigmoidal equation except for panel d where no sigmoidal fit was possible, a linear trend was used instead. *P*-values for Spearman rank correlations coefficients were computed with the following alternative hypothesis: Mean GI increases as K_d_s decrease. Correlations for unbinned data are reported in Extended Data Table 3. **e,** A section of the yeast GI network matrix, generated experimentally by synthetic genetic array (SGA). **f**, A reconstructed GI network matrix, generated computationally from a PPI network weighted by K_d_s. Hierarchical clustering was performed on the reconstructed matrix, and the resulting order of genes was applied to the experimental matrix. Full matrices are available on Figshare^45,46^.

The shapes of the curves are important in that they take the same forms as single site binding curves, with a flat lower end, dynamic range covering about two orders of magnitude, and flat saturation range. We speculate that these curves may represent an effective average binding curve with, for negative GIs, centered around average K_d_s of 1 μM and 10 μM for mouse and human, respectively and mean GI scores for both species at about −1.4. The curve for positive GIs in yeast is notable in that the half-maximal Mean GI Score is shifted to higher affinity (0.1 μM), consistent with the postulate that positive GIs are stronger for higher affinity PPIs^44^.

The strong correlation of mean PPI K_d_ and negative GI suggests that PPI binding affinity is a major driver of negative epistasis between the genes that code for the interacting proteins. These results imply that one network could be predicted from the other. To test this idea, we devised an approach to reconstruct GI network matrices from binding data alone (see Methods, Extended Data Fig. 6). We observed that the clusters along the diagonal of a matrix reconstructed from PPI binding data in yeast qualitatively matches that of experimental results in genetic screens (Fig. 2e, f)^1^. Overall, these results indicate that there exists a strong relationship between the quantitative features of GI and PPI networks, and that the modular structure of quantitative PPI networks seem to match that observed in GI networks.

### Redundancy and divergence of paralogs are reflected in genetic epistasis and PPI affinities

In our results above, we hypothesized that PPI, specifically binding affinity (K_d_), is an accurate proxy for cellular function and thus, it quantitatively reflects epistasis across the genome. Since the number of protein complexes is a function of abundance and K_d_, this poses a specific prediction that genomic events resulting in a maintained decrease or increase in abundance over macro-evolutionary time scales will be accompanied by a change in K_d_. In particular, gene duplication events, which could effectively double the number of cellular copies of a protein, could relax selection on K_d_ and still maintain the same number of complexes. Correspondingly, due to the relaxed selection on K_d_, we also expect a reduction in genetic interaction.

To test this hypothesis, we identified paralogs in our dataset and grouped them into four distinct types: 1) PPIs in which neither protein has a paralog, 2) PPIs in which one of the two (but not both) proteins has a paralog, 3) PPIs in which both proteins have a paralog, and 4) PPIs in which a protein interacts with its own paralog (paralogous heteromers); (Fig. 3, legend). Consistent with our expectation and previous analyses of GIs in paralogs, duplicated genes show weaker GIs with other genes than non-duplicates genes (Fig. 3a)^47^. This effect seems to be additive overall, as gene pairs where both genes have paralogs show weaker GIs (Fig. 3a, green). Remarkably, a similar trend is observed for binding affinity, where proteins encoded by duplicated genes tend to form weaker (high K_d_) PPIs (Fig. 3b).

**Figure 3.**
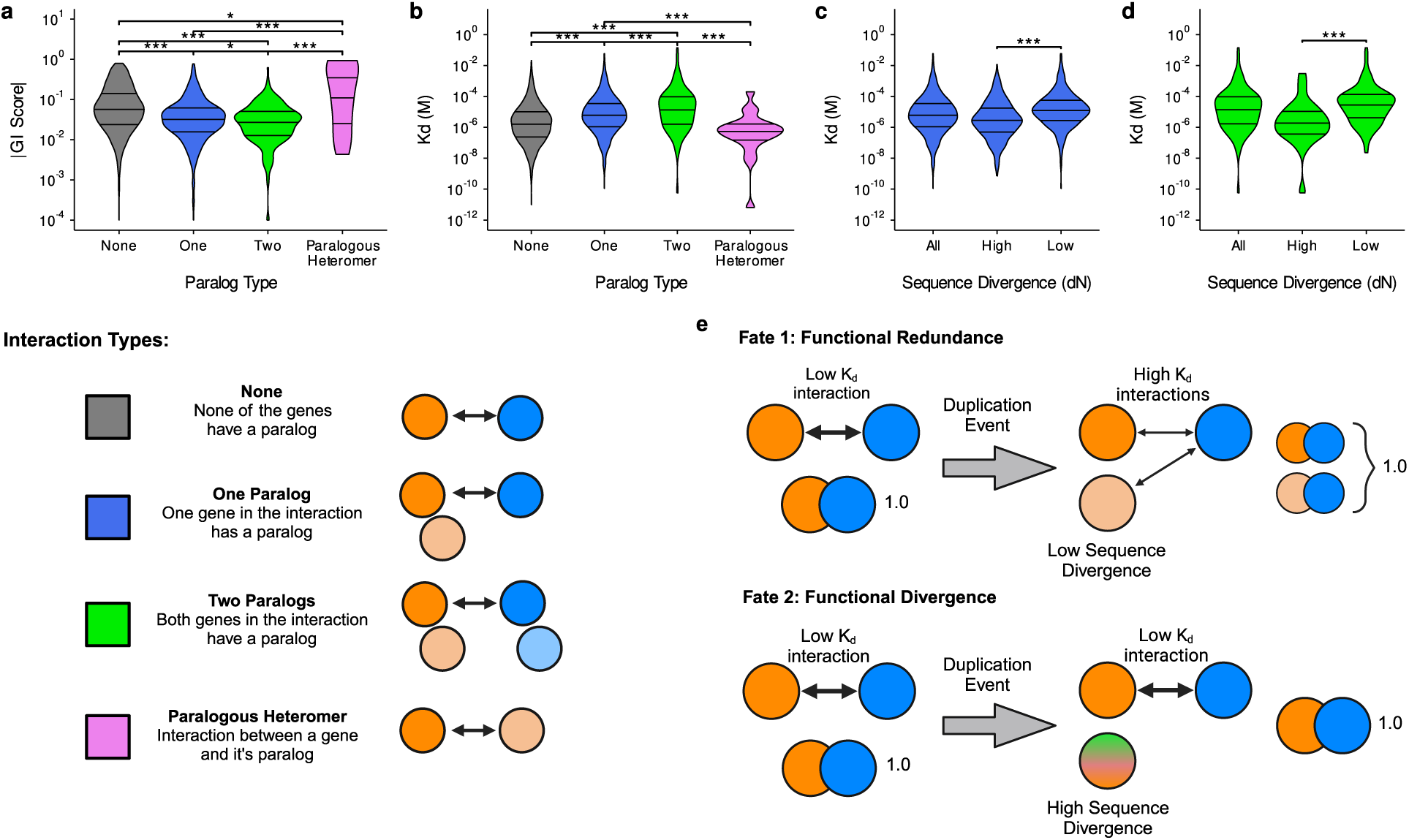
Functionally redundant paralogs participate in low binding affinity PPIs. **a,** GI scores for gene pairs which have one or two paralogs are lower than those without paralogs. Distributions for individual GI type (positive or negative) are included in Extended Data Fig. 7. **b,** Dissociation constants (K_d_) between proteins involved in a PPI are higher (lower affinity) if one or both proteins have a paralog. **c, d,** PPIs in which one (c) or both (d) proteins have a paralog separated according to sequence divergence. Significance levels are annotated as: * for *p* ≤ 0.05, ** for *p* ≤ 0.01, and *** for *p* ≤ 0.001. **e,** Hypothesized evolutionary model where duplicated genes with low sequence diverge (Fate 1) participate in weaker interactions over time. For duplicated genes with high sequence divergence (Fate 2), one paralog loses (or greatly reduces) their affinity for common interactors. Illustrations in legend and (e) were made with BioRender.

Intriguingly, two distinct modes are observable in the distribution of K_d_s for interactions in which both proteins have a paralog (Fig. 3b). This result is reminiscent of results from analyses of trigenic GIs involving duplicated genes, in which functionally redundant paralogs form more trigenic interactions (as opposed to digenic interactions) than functionally divergent paralogs, thereby resulting in a distinct bimodal distribution^48^. To test whether the separation of paralogs into redundant and divergent groups drives the bimodality observed in the K_d_s, we computed the sequence divergence (defined here as the number of non-synonymous mutations per non-synonymous site) between genes and their paralogs and separated them accordingly into “high” and “low” divergence paralogs (Extended Data Fig. 8, Supplementary Data 8). For interactions in which only one protein has a paralog, highly divergent paralogs participate in stronger interactions (lower K_d_) than paralogs with similar amino acid sequences (Fig. 3c); this is also true for interactions in which both proteins have a paralog (Fig. 3d).

It is known that some duplicated genes have conserved sequences and functions over evolutionary time^49,50^. However, protein abundance is expected to increase in the short term after gene duplication events, resulting in an imbalance of concentrations of protein complex components, which in turn negatively impacts fitness^51,52^. To reconcile functional conservation and concentration imbalance, it has been proposed that for short evolutionary time scales, post-translational expression attenuation is an important mechanism for robustness against an abrupt abundance increase following gene duplication and, over larger time scales, reduced expression allows for maintenance of redundant duplicated genes^53,54^.

However, reduced expression is not observed for all gene duplication events, suggesting other mechanisms may be at play^54^. In light of our results, we propose a model to explain how, on longer evolutionary time scales, protein complexes adapt to increased expression of their subcomponents after gene duplication (Fig. 3e). We hypothesize that following a duplication event, paralogous genes that conserve their sequence will conserve their interaction partners but participate in weaker interactions overall to balance the binding equilibrium such that the number of complexes stays the same (Fig. 3e, Fate 1). This can happen, for instance, when mutations occur in a few important amino acids at the binding interface or allosteric sites of paralogs, or if their interacting partners diverge such that their binding affinity for the duplicated genes is decreased. Furthermore, the subset of highly divergent paralogs participating in high affinity interactions (Fig. 3c and d) can be explained by a second scenario, where one of the duplicated genes diverges and loses (or drastically reduces the affinity for) interacting partners while the other maintains high affinity interactions (Fig. 3e, Fate 2). This model is both compatible with previous works showing that abundance is decreased following gene duplication^53,54^ and explains the tendency of highly divergent paralogs to make high affinity PPIs.

### Compositions of module and connector regions of the PPI network overlap with corresponding GI network regions

The classical interpretation of GI and PPI overlap mainly focuses on GIs within and between modules^4,7,19^; commonly referred to as the “within pathway” and “between pathway” models, respectively^17^. Indeed, protein-protein interaction modules (“complexes”) are reflected in the functional modules in GI networks^1,7,23^. However, not all proteins that participate in strong interactions form stable complexes of specific function; some participate in transient interactions that “connect” protein complexes together^55^. The degree to which these “connectors” in PPI networks are reflected in GI networks has not been explored. We hypothesize that the ability of certain proteins to connects complexes in PPI networks also holds for GI networks. In other words, if a protein connects two complexes (modules) in the PPI network, it will likely connect the same two functional modules in GI networks.

At the cellular level, functional modules involving proteins are typically metabolic pathways, signaling pathways or protein complexes^56^. However, we specifically limit our analysis of protein modules to solely include stable protein complexes in yeast, as defined by the ComplexPortal database^57^. Furthermore, GI modules were taken from an analysis of the double knockout GI network used in the previous sections^1^. For both GIs and PPIs, connectors were defined as the set of genes or proteins that did not fall into a module. As a preliminary step, we sought to determine the extent to which proteins/genes in modules overlapped between the two networks using this classification. We observe a significant overlap between the modules and for connectors for both networks *(p*-value = 1×10^−49^, hypergeometric test) (Fig. 4b, left panels).

**Figure 4.**
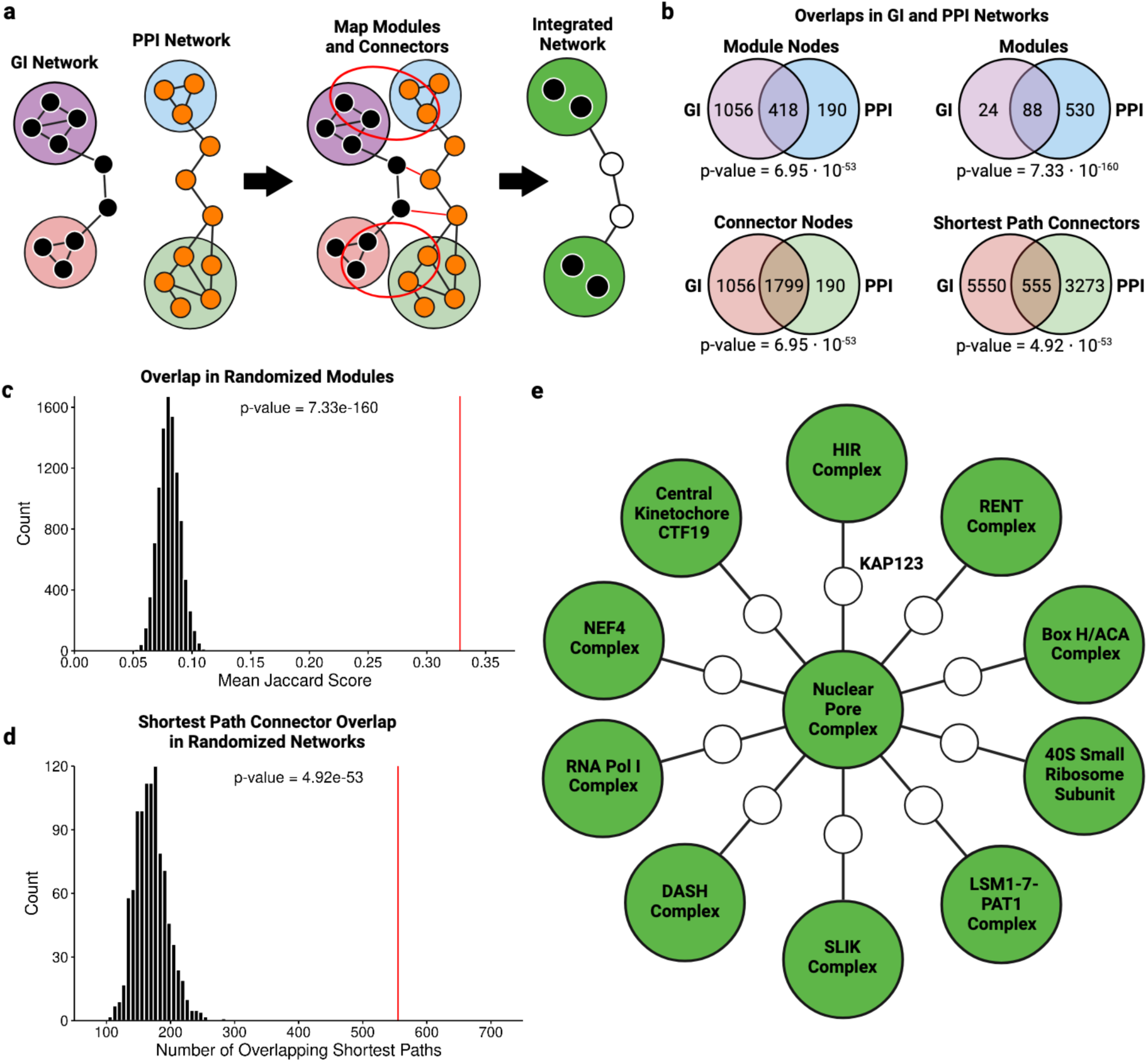
Integrating genetic interactions networks and protein-protein interaction networks using common topological structures. **a**, Brief description of the algorithm devised to identify connectors of interest. First, shortest paths between modules in GI and PPI networks are determined. Modules of GI and PPI networks are mapped 1:1 and their common nodes are identified; common shortest path nodes are also identified. The final integrated network is comprised of common components in the two networks. **b**, Summary statistics of different overlapping components in GI and PPI networks. **c**, A statistical validation of overlaps between modules in GI and PPI networks by comparing the observed overlap with that of randomized modules. **d**, A statistical validation of overlaps between shortest path in GI and PPI networks by comparing the number of observed shortest path overlaps to that of randomized networks. **e**, A subset of the integrated network centered around the nuclear pore complex and the karyopherin KAP123. Illustrations in a, b and d were made using BioRender.

#### GI modules map directly to PPI modules

We next sought to determine whether GI modules could be mapped one-to-one with protein complexes. To this end, we evaluated the similarity in gene composition of all possible GI module and PPI module pairings and identified module pairs with the highest degree of overlap. Strikingly, of the 112 GI modules, 88 could be mapped to a protein complex (Fig. 4b, upper left panel, Supplementary Data 9), which is significantly more than would be expected by chance (hypergeometric test *p*-value = 7×10^−160^, Fig. 4c). This analysis is complementary to that of Costanzo et al.^1^, in that we use a set of manually curated complexes, as opposed to GO enrichment, to demonstrate correspondence between GI and PPI modules. The remaining GI modules are likely attributable to pathways not involving stable complexes. For instance, a GI module composed of ERG13 and MVD1 was determined to be enriched for steroid and isoprenoid synthesis but was not mapped to any known complex. It is also important to note that the large fraction of non-overlapping protein complexes reflect the redundancy inherent to the ComplexPortal database, in which several variants or subunits of a complex are included as separate entries.

#### Connector proteins in PPI networks are analogous to connector genes in GI networks

The ability of connector proteins to bridge protein complexes has been highlighted in recent mass spectrometry screens of protein-protein interactions in human cells^20,21,55^. We reasoned that if most GI modules overlap strongly with protein complexes, some GI connectors may have topological properties similar to that of PPI connectors. The ability to bridge protein complexes being the most salient topological property of protein connectors, we focused on systematically identifying this motif network-wide for both GI and PPIs. Towards this aim, we devised an algorithm consisting of four main steps as illustrated in Fig. 4a for a single pair of modules from each network: 1) Mapping GI and PPI modules 1:1 as described above, 2) Finding the shortest paths between all pairs of modules; this was done independently in GI and PPI networks, 3) Identifying connector nodes (genes or proteins) in the shortest paths that are common between both networks, and 4) combining the common components into an integrated network. We identified 555 shortest path connectors common to both networks (Fig. 4b, lower right panel, Supplementary Data 10). Comparing this degree of overlap to that of randomized networks, we find that our results are extremely unlikely to arise by chance (hypergeometric test *p*-value = 4×10^−53^, Fig. 4d). Overall, our results demonstrate that connectors bridging modules are a topological feature shared by both GI networks and PPI networks.

#### Integrating PPI and GI networks highlights the functionality of connector genes and proteins

Previous analyses combining GIs and protein complexes have focused on GIs within and between complexes and the functional interpretations thereof^1,7,19^. Our approach to integrating PPI and GI networks into a “module-connector-module” structure could reveal novel functional relationships between the two. In fact, our integrated network reveals a core set of modules shared between both networks and highlights known functionality of well-studied connectors. For instance, a subnetwork of our integrated network correctly identifies the karyopherin kap123p as an important bridge between the nuclear pore and various complexes that are transported and/or localized in the nucleus (Fig. 4e). The integrated network also places RAS2 (Ras2p) adjacent to the adenylyl cyclase complex and the polarisome, which aligns with their respective roles in cell growth^58–61^. Interestingly, the integrated network also indicates that RAS2 connects the adenylyl cyclase complex to mitochondrial complexes, supporting a recently proposed mechanism in which Ras2p in the mitochondria promotes apoptosis via the cAMP/PKA pathway^62^. These examples highlight the potential of analyzing motifs in the connector region for generating new hypotheses on the functions of genes.

## Discussion

Determining how GI and PPI networks are related is an outstanding problem in systems and cell biology and its resolution has profound implications for our understanding of the genotype-phenotype relationship. While some relationship between binding affinity and GI strength could fit intuitive expectation, a quantitative relationship between binding affinity and the continuum of GI values has never been demonstrated. In this study, we examined the two properties that define biological networks (weights and topology) to establish a rigorous framework through which PPI and GI networks can be analyzed, predicted, and integrated. Although we focus on networks generated with data from the model organism *Saccharomyces cerevisiae*, we find that the principles established in yeast translate well to human networks.

The relationship between binding energies and GIs was further examined by estimating PPI dissociation constants proteome-wide, which we found to be strongly correlated with values from literature. These K_d_ estimations were used as a baseline to construct new models relating protein binding to genetic interactions. Indeed, binning PPIs according to binding information enabled both a reconstruction of a yeast GI network which qualitatively matched experimental results and the identification of an equation relating the general quantitative relationship between GIs and K_d_:

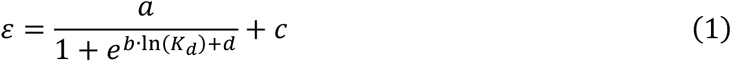

Where K_d_ is the dissociation constant of a PPI, ε is epistasis (multiplicative model) between the interacting proteins’ genes, and *a, b, c* and *d* are free parameters. Importantly, these parameters have tangible biological and biophysical meanings. For instance, the parameter *a* sets the range of epistasis values. Further, the parameter *b* (replaced by kT below), the parameter *d*, and K_d_ form the Gibbs free energy of binding equation:

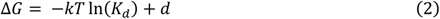

We note that equation 1, relating K_d_ and epistasis is a sigmoidal function, having the same form as the equation relating K_d_ and fraction bound. The implication of this correspondence between GI and PPI networks is profound: Tracing non-additive effects (epistasis) to their molecular thermodynamic underpinnings implies the possibility of modelling effects of individual genetic variations on a given trait (genotype-to-phenotype relationship). A caveat of our analysis is that epistasis is defined specifically in the context of pairwise gene deletions (or temperature sensitive mutant gene knock downs) as opposed to other type of genetic changes such as point mutations, recombination, or structural variants. Moreover, our model for the reconstruction of a complete GI network (Fig. 2f) explains approximately 2% of the entire variance in the negative GI network, including GIs between genes whose proteins do not interact with each other. Although a better understanding of how genetic perturbations percolate through the PPI network would likely increase explainability, it is likely that GIs are also strongly driven by other molecular mechanisms such as substrate and product dependencies between enzymes in metabolic networks, and transcription factor-promoter relations in gene regulatory networks.

Certain limitations exist as well regarding our estimates of dissociation constants. First, each PPI is treated as completely independent of any other, leading to deviations from the actual values. For instance, competitive binding will systematically lead to underestimations of binding strength, especially for very strong interactions. Second, under our current model, interaction stoichiometries over 1:1, which may occur if more than one prey protein binds to a single bait protein, lead to negative K_d_ values which are physically impossible. The number of interactions which are discarded as a result represents a non-negligible fraction of the interactome, which may bias the distribution of our estimated K_d_. Third, our ability to assess the precision of our estimates is limited to the quality of reference binding affinity data in the literature. Fourth, we assume all reported interactions in our datasets to be direct. In reality, AP-MS approaches also detect indirect interactions which are not readily distinguishable from direct ones.

Previous approaches aiming to identify common topological features between GI networks and PPI networks have mostly focused on GIs within or between protein modules^4,7,19^. However, not all genes and proteins cluster into highly connected groups. Although it has long been known that certain proteins serve as “connectors” between protein modules^63^, the extent to which this “module-connector-module” motif occurs in GI networks had not been explored. We demonstrate that the common topological organization in GI and PPI networks extends further than overlap in modules, but also the connectors that bridge these modules. By merging modules and connectors from both GI and PPI networks an integrated network was constructed, where the functional role of connector genes/proteins are highlighted and can be used a novel tool for functional inference that will be valuable for predicting GIs in other organisms.

Finally, the quantitative and topological relationships we have established in this study could provide means of interpreting and predicting fundamental relationships between genes underlying normal and disease phenotypes, as well as identify specific vulnerabilities or changes in PPIs that could predict potential therapeutic targets for diseases.

## Methods

### Collecting yeast and human genetic and protein-protein interaction data

For yeast, raw synthetic genetic array (SGA) genetic interaction data was downloaded from TheCellMap^1,64^ in matrix format. Data from essential and non-essential strain crosses (Essential vs. Essential, Essential vs. Nonessential, Nonessential vs. Nonessential) were combined. In instances where multiple GIs were measured for a pair of genes, only the entry with the most significant *P*-value was retained. For human cell lines, GI data was retrieved from a digenic knockdown screen of 472×472 genes in Jurkat and K562 cells^35^ (table S5 of reference 35). For both yeast and human data, unless otherwise indicated, analyses were performed using all GIs regardless of significance scores.

Yeast protein-protein interactions were taken from a near exhaustive whole-proteome affinity enrichment coupled to an affinity purification-mass spectrometry (AP-MS) screen^22^ (Yeast_Interactome_Edges.csv from supplementary file 2 of reference 22). Human protein-protein interactions and stoichiometries were taken from two quantitative AP-MS screens: one in which genes were tagged with GFP and expressed on bacterial artificial chromosomes in HeLa cells^20^ (table S2 in reference 20), and a second in which genes are endogenously tagged in HEK 293T cells with the small fragment of the split mNeonGreen_2_ tag^21^ (table S4 in reference 21). Both human cell line interactomes were concatenated but common interactions between the two screens were kept as separate data points for all analyses.

### Computing interaction stoichiometries and abundance stoichiometries in yeast

Interaction stoichiometries were calculated from data generated in an AP-MS screen^22^ using a previously described strategy^20^. The starting preprocessed table containing normalized and imputed label-free quantification (LFQ) intensities from the AP-MS screen were provided by the authors. Briefly, computing interaction stoichiometries was done in three mains steps: 1) Subtracting the median intensity of a protein across all sample, 2) Dividing LFQ intensities by the number of theoretically observable peptides of lengths between 7 and 30 amino acids, 3) dividing the values observed for the prey protein by that of the bait protein. Importantly, the number of theoretically observed peptides were computed using sequences from the Uniprot database^65^ (https://www.uniprot.org/proteomes/UP000002311) using LysC only digestion (as opposed to LysC and Tryptic digestion). Negative interaction stoichiometries (which arise because of normalization in step 1) were dropped from the dataset. The final dataset includes interaction stoichiometries for 7348 unique PPIs.

Protein abundances were taken from a dataset unifying measurements from various sources^30^ (table S4 in reference 30). Abundance stoichiometries were determined by calculating the ratio between the median copy numbers of the prey over that of the bait, as described previously^20^. Abundance measurements for both interactors were available for 7336 out of the 7348 PPIs for which interaction stoichiometry values were calculated. The 22 interactions which had interaction stoichiometry values, but not abundances stoichiometry values, were conserved in the dataset for analyses in which abundance stoichiometry was not relevant.

### Analysis of genetic interactions and protein-protein interaction stoichiometries

The relationship between GIs and stoichiometries was explored in yeast and human cell lines (see first section for data collection). To merge GI and PPI stoichiometry datasets, tables were merged such that if genes comprising a GI also code for proteins that interact, the GI score is assigned to that PPI. Four stoichiometric regions were graphically defined within stoichiometry plots according to previous criteria^20^. Hein et al. defined region 1 as the space enclosed by a circle centered at (−0.5, 0) where the abscissa is interaction stoichiometry, the ordinate is abundance stoichiometry, and coordinates are in base 10 logarithmic scale. Using these definitions as a reference, region 3 was defined using circles of identical radii (0.75) centered at (−0.5, 1.5), region 2 was centered at (−2.5, 0) for yeast and (−3.5, 0) for human, and region 4 was centered at (−2.5, −1.5) for yeast and (−3.5, −1.5) for human.

### Estimating protein-protein interaction dissociation constants

To estimate dissociation constants from quantitative mass spectrometry data, we first assumed all individual PPIs to be independent of all others (no competitive binding) and their binding dynamics to be describable by a two-state system:

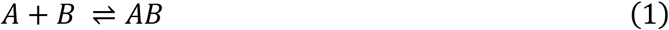

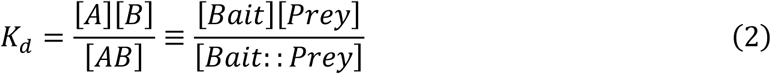

Where K_d_ is the dissociation constant, [A] the molar concentration of unbound protein A, [B] the concentration of unbound protein B, and [AB] the concentration of the complex formed by proteins A and B. In a given mass spectrometry experiment, one of the two proteins will be the bait (tagged protein retained on column) and the other a prey.

Next, we sought to estimate the number of bait-prey complexes using interaction stoichiometries. By definition, interaction stoichiometry is a proxy measurement of the ratio of prey protein bound to bait protein over the total abundance of bait protein. Under the previously mentioned assumptions, the amount of prey bound to bait is equal to the number of bait-prey complexes.

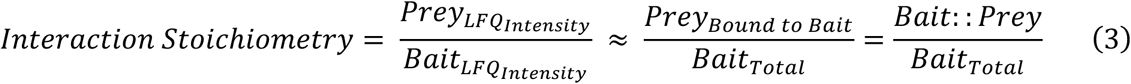

Therefore, we used the product of interaction stoichiometry and total abundance of bait (in copy numbers) to estimate the number of bait-prey complexes in a given cell.

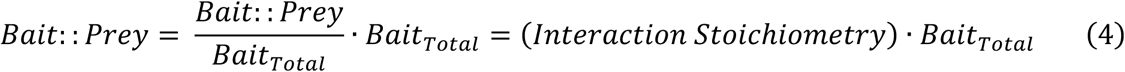

For yeast PPIs, the cellular copy numbers of bait proteins were taken from the same source as that described to calculate abundance stoichiometries^30^. For PPIs in human cell lines, abundances were taken from whole proteome quantification experiments which were performed alongside the AP-MS measurements of interaction stoichiometries^20,21^ (opencell.czbiohub.org/data/datasets/opencell-protein-abundance.csv [downloaded April 4^th^ 2022]; Hein et al., table S3).

Next, the number of unbound bait and prey proteins were estimated by subtracting the number of bait-prey complexes^40,66^. In a few instances, the estimated number of bait-prey complexes was higher than that of the total abundance of bait or prey, resulting in a negative number of unbound proteins. This almost only arose when interaction stoichiometry is above 1. These negative values were removed.

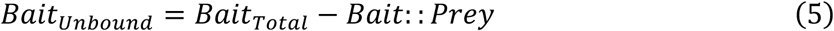

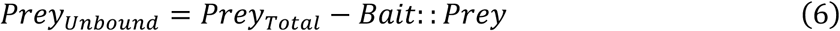

Copy numbers of bait proteins, prey proteins and bait-prey complexes were next converted to molar concentrations (mol/L) using Avogadro’s constant and cell volumes. Below is an example formula for bait concentrations; concentrations prey and bait-prey complexes were calculated in the same way. Dissociation constants were calculated using the resulting concentrations.

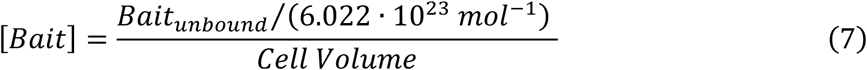

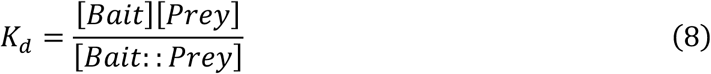

A cell volume of 4.2 × 10^−14^ Liters was used for yeast PPIs (which were measured in haploid cells)^67^. A cell volume of 2.6 × 10^−14^ Liters was used for PPIs measured in HeLa cells by Hein et al.^68^ Cho et al. measured PPIs and stoichiometries in HEK293 cells, for which no experimental measurement of volume was found. However, the diameter of HEK293 cells was measured to be approximately 13 micrometers (Bionumbers ID: 108893). Therefore, an estimated cell volume of 1.15 × 10^−14^ Liters was calculated from this diameter by approximating the shape of a cell to that of a sphere. The above-mentioned values were all found via the BioNumbers database^69^.

### Validating estimated dissociation constants using values from literature

Experimental dissociation constants from literature were taken from the protein-protein complex subset of the PDBbind database (2020 version)^70,71^. Taxonomic identification numbers were extracted from the PDB file of each database entry. Only interactions involving proteins from budding yeast and human were retained. PDB files which listed more than two proteins were removed, as it would not be possible to assign dissociation constants to the correct protein pair.

Systematic names or IDs for proteins and genes are absent from the PDBbind database. For this reason, the PDB ID for each complex was used to access their structure’s RCSB Protein Data Bank web page and retrieve the Uniprot IDs of the proteins within the structure. Web scraping for the data published in this article was performed on April 23^rd^ 2024, using the *BeautifulSoup4 Python3* package. For yeast PPIs, Uniprot IDs were then converted to systematic ORF IDs using the *org.Sc.sgd.db R* package. For human PPIs, a Uniprot to HGNC symbol conversion table was downloaded (on February 28^th^ 2024) from the “Custom downloads” page of the HGNC website^72^. Information on the original source, which is often different from that of the structure, and details on the experimental setup of each dissociation constant measurement is also absent from the database. Although this limits quality control, it is known from curated sets for structure-based modelling of PPI affinities that common methods to measure PPI binding affinities are best for interactions in the micromolar to nanomolar range^42^. Therefore, measurements were limited to dissociation constants no smaller than 1 nM and no greater than 1 mM (1 nM ≤ K_d_ < 1 mM). The final reference dataset contains 505 K_d_ measurements for human PPIs and 59 for yeast PPIs.

Estimated K_d_ values for PPIs were merged with experimental values for the reference dataset. For PPIs in human cell lines, 70 estimated K_d_ values were mapped to values from literature. Only 4 estimated K_d_ values were mapped to values from literature for yeast PPI.

### Comparison of mass spectrometry K_d_ estimates in yeast using PCA K_d_ estimates

Colony intensities and z-scores were collected from a genome-wide screen for protein-protein interaction using a survival-selection protein-fragment complementation assay (PCA)^73^ (table S4 of the referenced article) and filtered to only contain the final 2770 protein-protein interactions (table S3 of the referenced article) reported in the article. Next colony intensities were collected from a proteome-wide abundance screen based on the GFP-PCA method^74^ (Data Set S2 of the referenced article). GFP-PCA intensities were merged to cellular copy numbers from a compilation of abundances in yeast^30^ to generate a linear regression model relating PCA colony intensities to copy numbers. Protein-protein interaction PCA intensities were then rescaled to a range similar to that of the abundance GFP-PCA intensities. The linear regression model was then used to convert protein-protein interaction intensities to numbers of complexes. With the number of copies of complexes and individual protein copy numbers at hand, dissociation constants were estimated as described in the “Estimating protein-protein interaction dissociation constants” section. In total, 1480 K_d_ estimates were calculated from PCA data, of which 48 also had K_d_ estimated from mass spectrometry data.

### Reconstructing a GI network matrix from a PPI binding network

Yeast GI and PPI dissociation constant (K_d_) tables were merged such that if genes comprising a GI also code for proteins that interact, the GI score is assigned to that PPI. The final table contains 3265 interactions, each with a GI score and a K_d_ value. A graph was then generated, using individual proteins as nodes/vertices (*V_PPI_*), PPIs as edges (*E_PPI_*), and the reciprocal negative logarithm of K_d_ values as weights (*W_PPI_*):

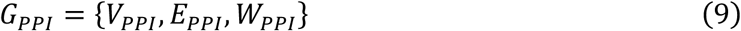

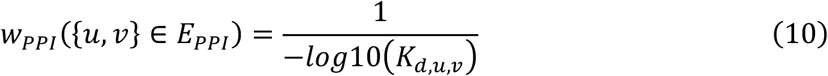

Where *u, v* ∈ V_PPI_, K*_d,u,v_* is the dissociation constant between the interacting proteins *u* and *v*, and *w_PPI_*(*e*) is the weight assigned to the edge *e* (where *e* = {*u*, *v*} connects *u* and *v*).

To establish an equation relating weights and GI scores, PPIs were first binned into quantiles according to their weights. The mean weight and mean GI score within each quantile were then calculated and an exponential equation was used to fit the data (See Extended Data Fig. 6a for example with 25 quantiles). The root mean square error (RMSE) of the fit for different numbers of quantiles were tested to find the most optimal (Extended Data Fig. 6b). A dip of RMSE values and narrowing of RMSE spread was observable at around 25 quantiles, which justified its choice as the ideal number of quantiles for the final fitting (Extended Data Fig. 6a, c).

To mimic the completeness of GI networks, the PPI graph was converted to an “association” matrix similar to that described by Rives and Galitsky^75^. Briefly, the *shortest_path_length* function of the *Networkx* python library was used on *G_PPI_*, using *W_PPI_* as the weight parameter, to generate a |*V*| × |*V*| matrix (*d*) in which d[i,j] denotes the length of the shortest path between the nodes (proteins) i and j. This all-pairs-shortest-path (APSP) matrix was used as a graph (*G_APSP_*), such that:

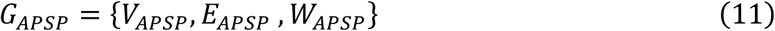

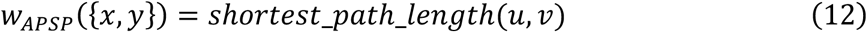

Where {*u*, *v*} ∈ *E_PPI_*, {*x*, *y*} ∈ *E_APSP_*, and both edges represent the same protein interaction. The exponential equation relating means of quantized weights and GI scores was then used on *W_APSP_* to generate a new graph (G*_GI_pred_*) with weights *W_GI_* containing estimated GI scores. The adjacency matrix of G_GI_pred_ was clustered with average linkage hierarchical clustering using Pearson correlation in Cluster 3.0 (*74*).

### Analysis of dissociation constants in yeast paralogs

Whole genome duplication (WGD) paralogs were collected from Kellis et al.^76^ (Supplementary file 8 of the cited article) and small-scale duplication (SSD) paralogs were collected form VanderSluis et al.^47^ Paralogs were separated into 3 categories as described in the main text. The rate of non-synonymous mutations per non-synonymous site (dN) was used as a proxy for sequence divergence. To calculate dN values, paralog pairs were aligned pairwise at the protein level with ClustalW (version 2.1) using the PAM protein weight matrix^77^. Using the *reverse.align* function of the *seqinr* R package, protein-coding nucleic acid sequences were next aligned using protein alignments as a guide^78^. The reverse alignments of each paralog pair were then used to compute dN using the *kaks* function of *seqinr*. Amino acid sequences and coding nucleic acid sequences were downloaded from the SGD sequence archives (http://sgd-archive.yeastgenome.org/sequence/S288C_reference/, March 2024) for the S288C reference strain. A sequence divergence threshold of 0.2 was selected to categorize paralog pairs into “high” and “low” sequence divergence (dN) (Extended Data Fig. 8a). For protein-protein interactions where both interactors have a paralog, the interaction was categorized into “high” when both interactors had a dN value above 0.2.

### Mapping GI Modules to PPI Modules

GI clusters were downloaded from an analysis of a near-complete double knockout study in yeast^1^ (Data File S6 of the cited article). Yeast protein complexes were downloaded from a species-specific dataset of the ComplexPortal database^57^; protein Uniprot IDs were converted to systematic yeast ORFs using the *org.Sc.sgd.db* R library^79^. All GI clusters were compared to all PPI complexes by their common ORFs, the degree of overlap was scored using the Jaccard index. For each GI cluster, the protein complex which had the highest overlap (Jaccard Index) was retained.

If a GI cluster overlapped better with connector nodes (proteins outside complexes), it was dropped from the dataset. To validate the significance of the overlap, GI clusters were randomized and mapped to protein complexes. GI clusters were randomized by retaining the size of the clusters and randomly redistributing ORFs. The randomization was performed for 10 000 iterations on a general-purpose computer cluster. A p-value was derived from a z-score in which the mean Jaccard score from the experimental dataset was compared to the distribution of mean Jaccard scores of randomized clusters (Fig. 4c).

### Identifying Nodes Connecting GI and PPI Modules

The GI and PPI networks used for the analysis of “connector” nodes were taken from the near-complete double knockout screen in yeast and whole-proteome affinity enrichment coupled to mass spectrometry screen described in the *Collecting yeast and human genetic and protein-protein interaction data* section. Only negative genetic interactions were used for the analysis in Fig. 4. Because shortest paths are sensitive to noise, we used the moderately stringent cutoff of GI score < −0.08 and p-value < 0.05 recommended by the authors^1^. Identifying nodes (gene/proteins) connecting modules in GI and PPI networks was performed in 4 main steps. First, genetic modules and protein modules are mapped 1-to-1 as described in the *Mapping GI Modules to PPI Modules* section above.

The 1-to-1 mapping can be expressed mathematically, where *M_GI_* and *M_PPI_* are each a set containing all modules in the GI and PPI network, respectively, and a module is itself a set of genes/proteins. Connectors can be defined as the sets *C_GI_* and *C_PPI_*, which contain all the connector genes/proteins in the GI and PPI networks respectively. Importantly, a gene/protein cannot be both a connector and within a module. In other words:

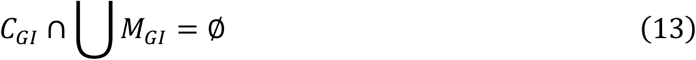

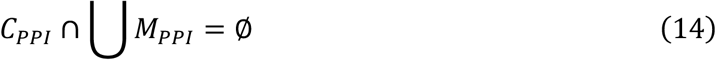

We note that here the large union symbol denotes the union of all genes/proteins that are within modules into a single large set. Next, a module *X* ∈ *M_GI_* is related (mapped) to a module *A* ∈ *M_PPI_* if and only if J(*X*, *A*) > J(*X*, *C_PPI_*), and J(*X*, *A*) > J(*X*, *B*) for all *B* ∈ *M_PPI_*. This relation (mapping) is denoted as *R*, where *(X, A)* ∈ *R* for all modules *X* in *M_GI_* and for all modules *A* in *M_PPI_*. Note that J denotes the Jaccard index:

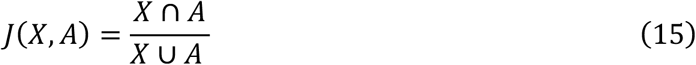

Second, for all combinations of modules within a network, the shortest path between all pairs of nodes were computed. This was done independently for GI and PPI networks. In mathematical terms, for a graph *G =* {*V, E, W*} (representing the GI or PPI network), let {*a, b, c*} be a set of genes within module *A* such that *A* = {*a, b, c*} and {*a, b, c*} ⊂ of *V*, and let {*x, y, z*} be a set of genes within module *B* such that *B* = {*x, y, z*} and {*x, y, z*} ⊂ *V*. Importantly, *B* **∩** *A* = ∅. For any pair (*u*, *v*) ∈ *A* × *B*, the shortest paths between nodes *u* and *v* were computed using Dijkstra’s algorithm (*all_short_paths* function of the *networkx* python package).

Third, connector nodes that bridge the same modules in both GI and PPI networks were systematically identified. If the sequence of nodes (*i, j, k*) is the shortest path between module *A* and module *B* in the PPI network (*G_PPI_*) and the sequence (*k, m, n*) is the shortest path between module *X* and module *Y* in the GI network (*G_GI_*), then (*i, j, k*) **∩** (*k, m, n*) is the shortest path common to both networks. Note that here modules *A* and *X*, as well as *B* and *Y* are mapped to each other; *(A, X)* ∈ *R* and *(B, Y)* ∈ *R*. Importantly, to avoid including nodes from the source and target modules in the shortest path, nodes belonging to a module were removed from the shortest path.

Fourth, the overlapping features in both networks *i.e.,* modules and shortest paths connectors, are combined to generate an integrated network. To determine if the extent of the overlap in connectors is due to network topology as opposed to random chance, the approach described above was performed on randomized networks. The networks were randomized by first conserving edges (interactions) within modules and then performing degree preserved randomization for all other edges. Overlapping connectors were identified for 1 000 random GI and PPI networks on a general-purpose computer cluster. A p-value was derived from a z-score in which the number of overlapping connectors from the experimental dataset was compared to the distribution of overlapping connectors from randomized networks (Fig. 4d).

## Acknowledgments

The authors thank Anastasia Baryshnikova for advice on preprocessing of synthetic genetic array genetic interaction data. We also thank Christian R. Landry, Ben Lehner, Elena Kuzmin, and Brenda Andrews for comments on the manuscript.

## Author contributions

SWM, AWRS and XCG conceptualized this study. XCG generated the computational pipeline. XCG, AWRS and SWM analyzed and interpreted data. XCG, AWRS and SWM wrote the manuscript.

## Data and materials availability

Full clustered Genetic Interaction Matrices are available on Figshare (*42-43*). All final and important intermediate data files are provided online and on GitHub (see link below). Original data sources are cited in Methods. Python, R and bash scripts are available in the following GitHub repository: https://github.com/Xavier-Castellanos-Girouard/PPI_Binding_Epistasis.

---

**Extended Data Figure 1.**
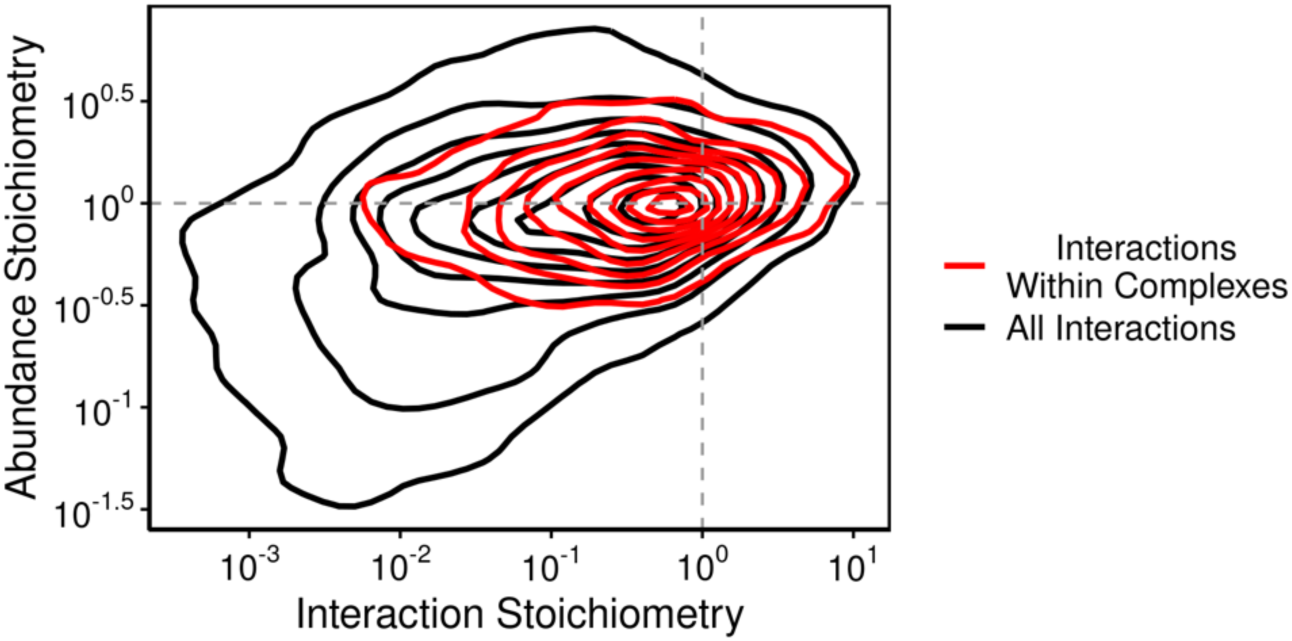
Stable protein complexes in *Saccharomyces cerevisiae* are enriched for 1:1 stoichiometries. Yeast PPIs occurring between proteins of stable complexes as annotated by the ComplexPortal database (*55*).

**Extended Data Figure 2.**
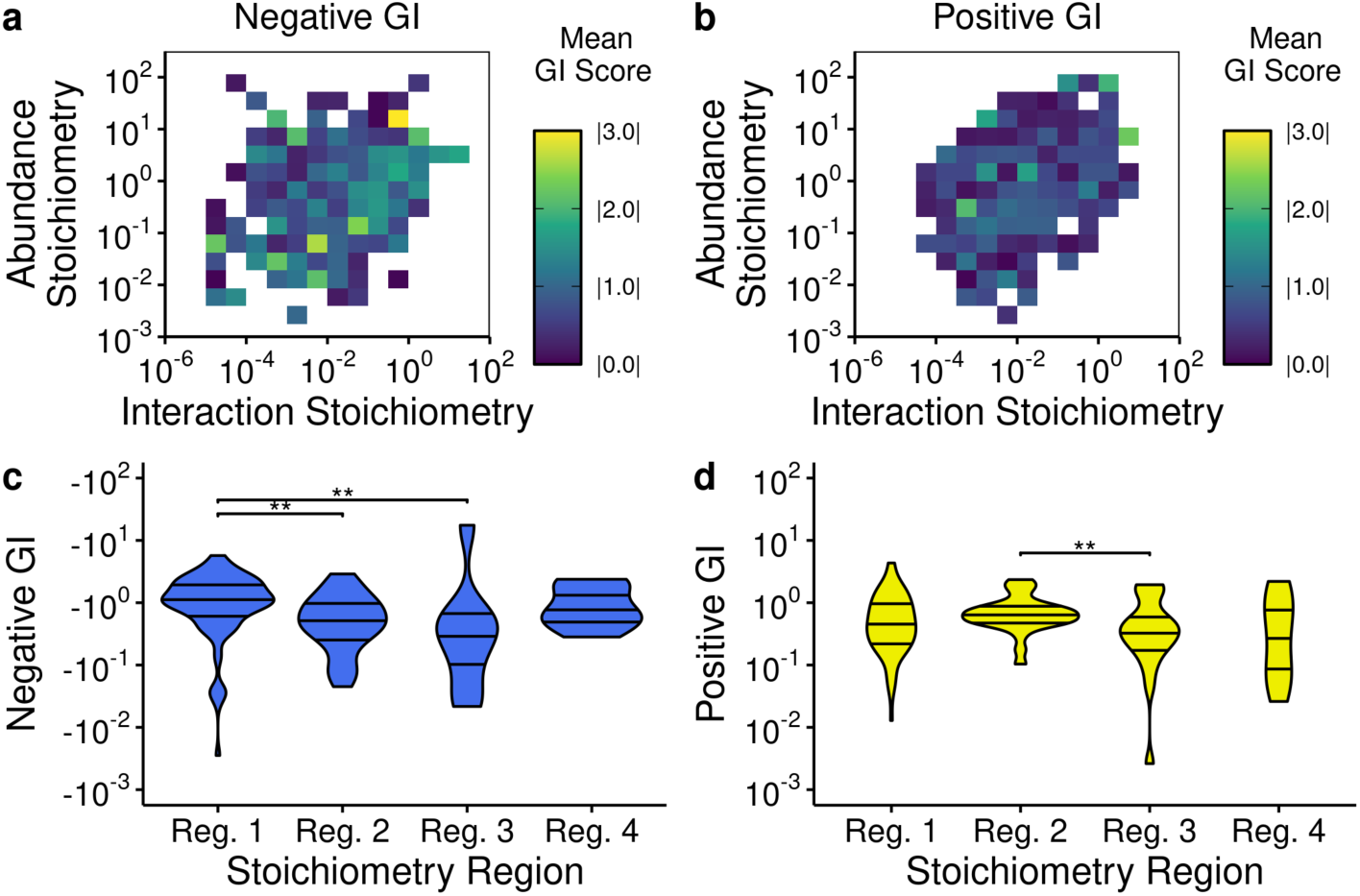
The PPI stoichiometry landscape of GIs in the Jurkat cell line. **a, b,** Mean negative and positive genetic interaction scores for PPIs throughout the stoichiometry plot. GI scores in human were taken from a screen in Jurkat cell lines. **c, d,** Distribution of negative (c) and positive (d) GI scores within the 4 defined regions of the stoichiometry plot. Significance levels are annotated as: ** for *p* ≤ 0.01.

**Extended Data Figure 3.**
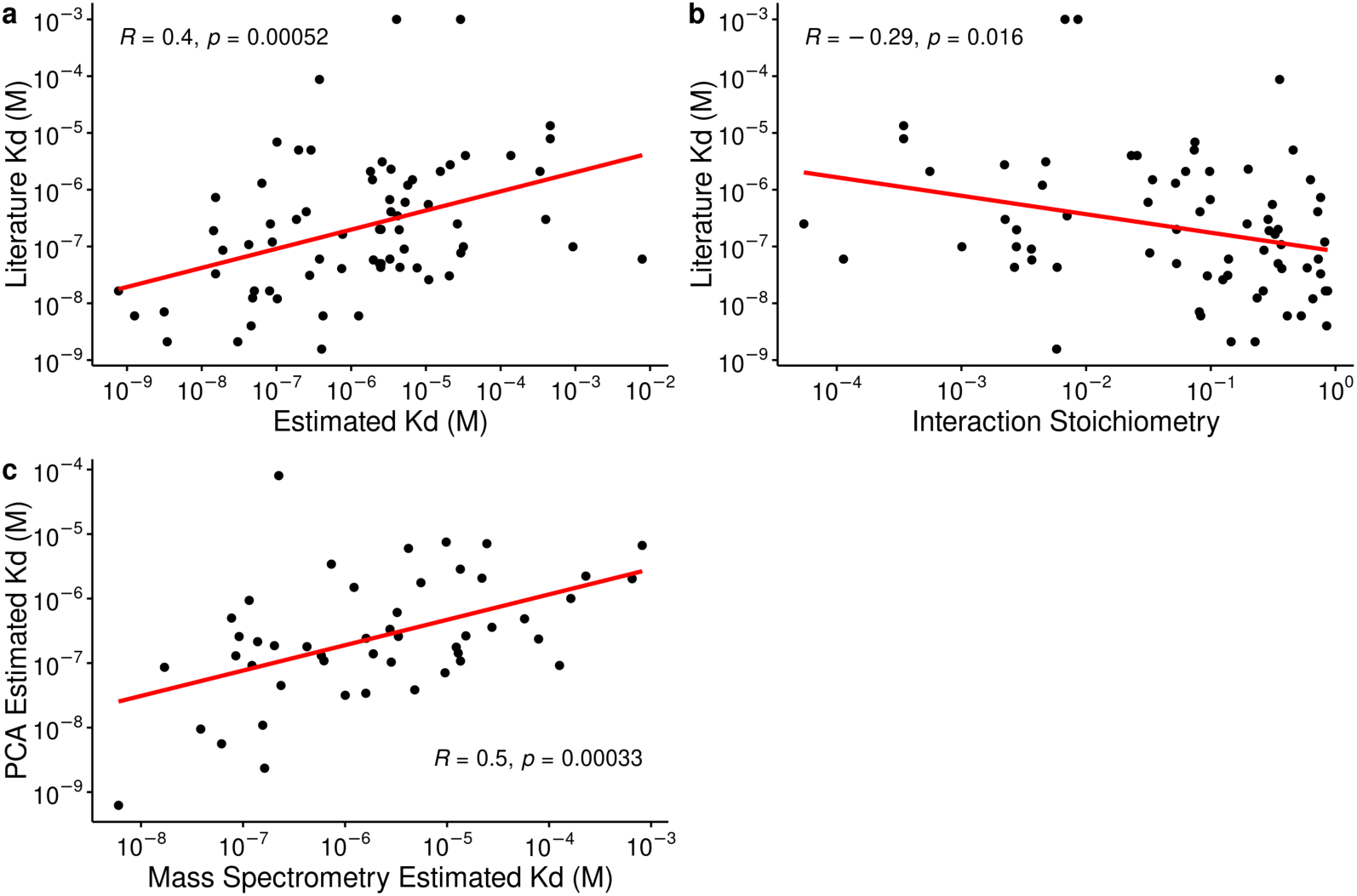
Validation of dissociation constants estimated from quantitative mass spectrometry measurements. Pearson correlation was used in all panels. **a,** Estimated PPI dissociation constants estimated from quantitative mass spectrometry data in human cell lines correlate moderately but very significantly with values from literature. **b,** Interaction stoichiometries from quantitative mass spectrometry data in human cell lines correlate weakly with values from literature. **c,** Dissociation constants estimated from quantitative mass spectrometry data in yeast correlate moderately but very significantly with those estimated from protein-fragment complementation (PCA) results.

**Extended Data Figure 4.**
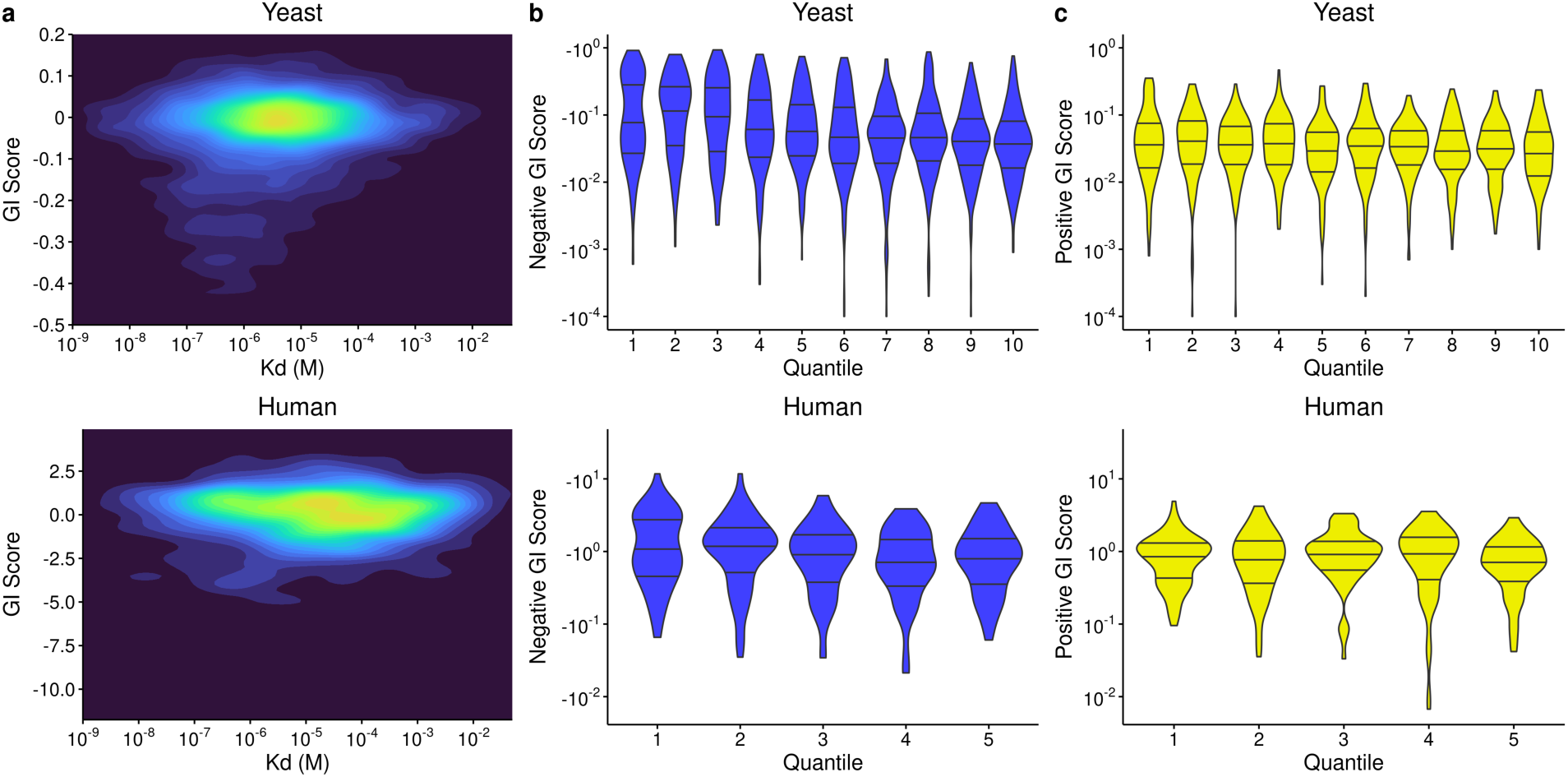
High affinity protein-protein interactions are biased towards strong GIs. **a,** Density plot of genetic interaction scores as a function of their protein-protein interaction affinity (K_d_) for yeast and human (K562 cell line). **b,** Distribution of negative GI scores across K_d_ quantiles (quantile 1 is the lowest K_d_; strongest affinity) for yeast and human. **c,** Distribution of positive GI scores across K_d_ quantiles (quantile 1 is the lowest K_d_) for yeast and human.

**Extended Data Figure 5.**
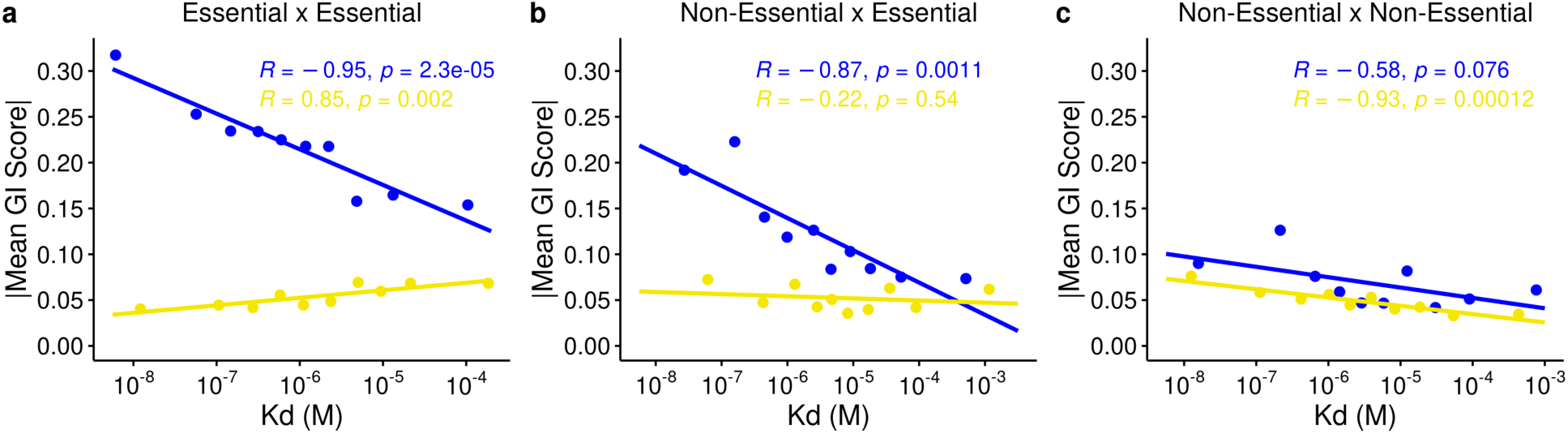
Correlations between binned genetic interactions and K_d_ for genes pairs of different essentialities. **a-c**, Pearson correlation coefficients between dissociation constants (K_d_) and GIs are computed for genes pairs where both genes are essential (a), only one gene is essential (b), or both genes are non-essential (c). *p*-values are computed from two tailed test. All trendlines are fit using a linear regression.

**Extended Data Figure 6.**
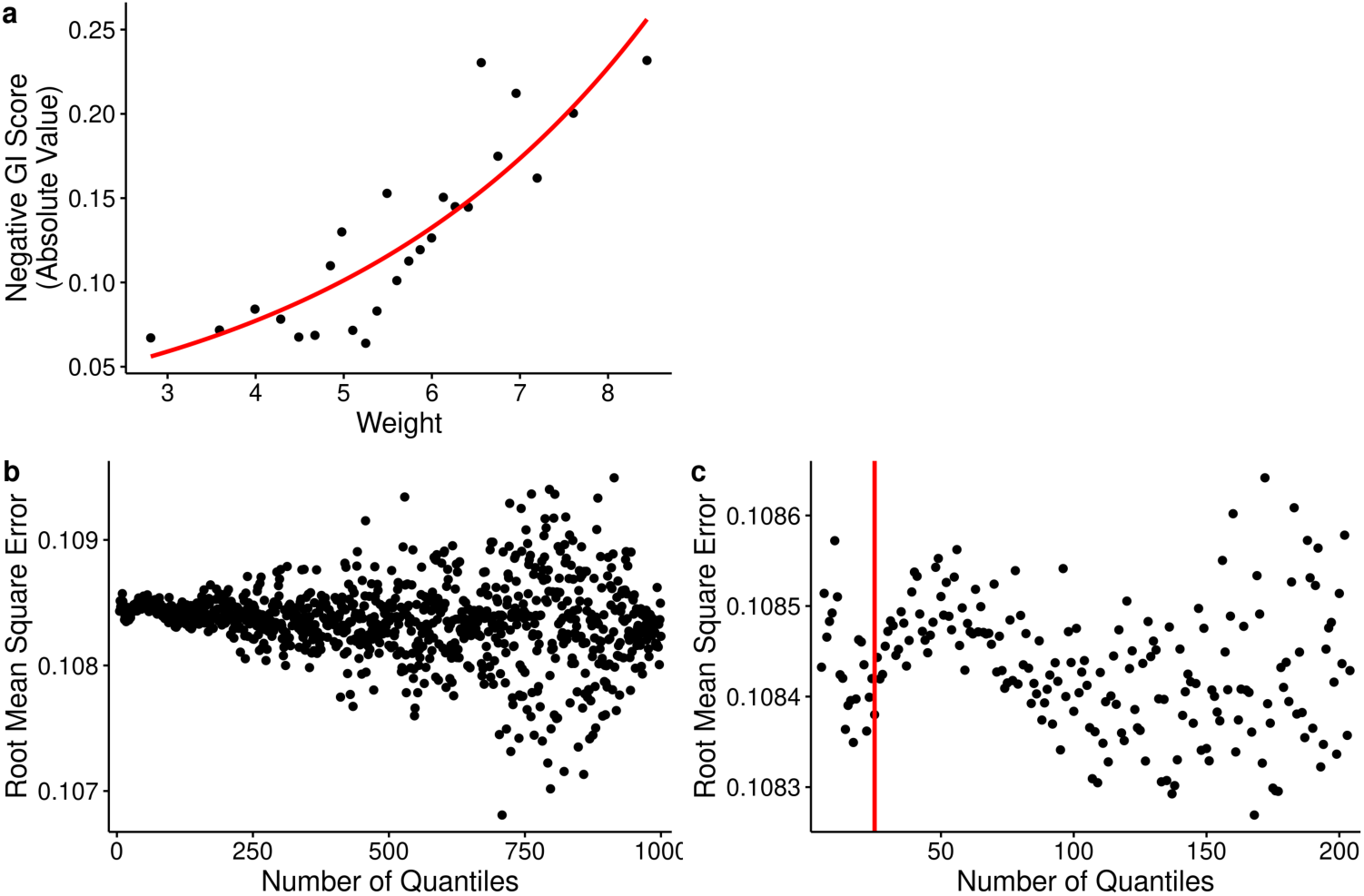
Fitting an equation relating binned PPI weights (derived from K_d_ values) to GI scores. **a,** Yeast PPIs are binned into 25 quantiles according to their network weights. The average weight and GI score is calculated for every quantile and an exponential equation is used to fit the data. **b,** The fitting procedure described in (a) was tested with different numbers of quantiles to visualize their root mean square errors (RMSE). **c,** A narrowing of RMSE spread and a dip in RMSE values can be observed at around 25 quantiles.

**Extended Data Figure 7.**
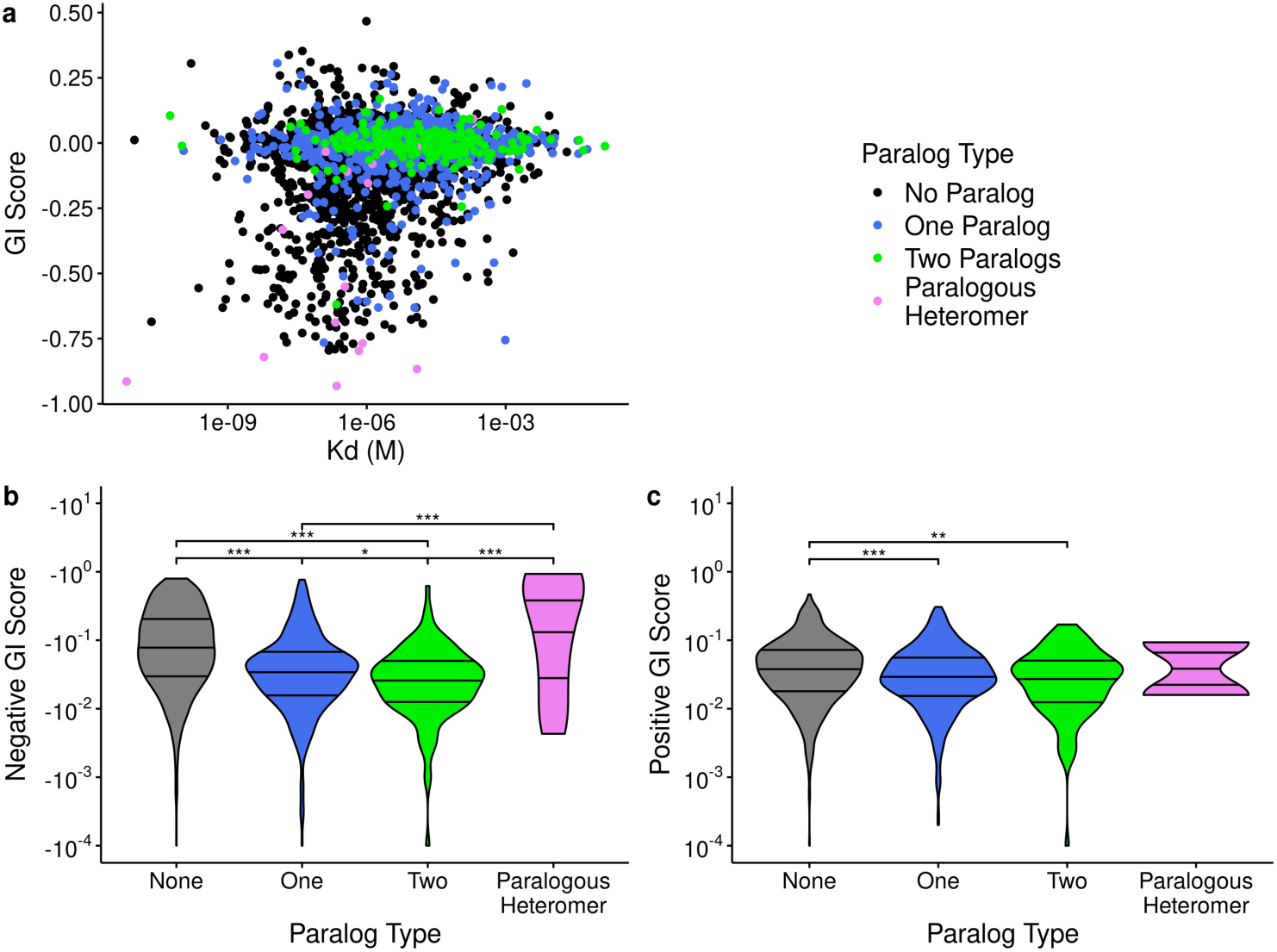
Distribution of positive and negative genetic interaction scores according to PPIs separated by their paralog composition. **a,** Distribution of estimated K_d_ and GIs (yeast dataset) according to the PPI’s paralog composition/type. **b,** Distribution of negative GI scores for different interaction type. **c,** Distribution of positive GI scores for different interaction type.

**Extended Data Figure 8.**
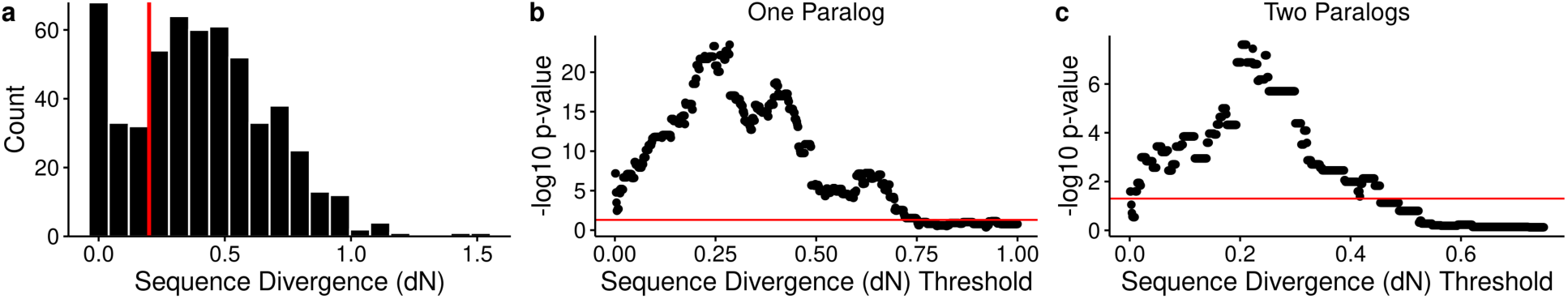
Dividing protein-protein interactions according to paralog sequence divergence. **a,** Sequence divergence (dN) between paralogous protein/gene pairs retrieved from literature. A dN threshold of 0.2 separates the two visible modes in the distribution. **b, c,** The statistically significant differences in dissociation constants between high divergence and low divergence are robust to the chosen dN threshold. This holds for PPIs in which one protein has a paralog (b) and PPIs in which both proteins have a paralog (c). *p*-values are calculated using Wilcoxon rank sum test. Red line indicates a *p*-value of 0.05.

**Extended Data Table 1^a^:**
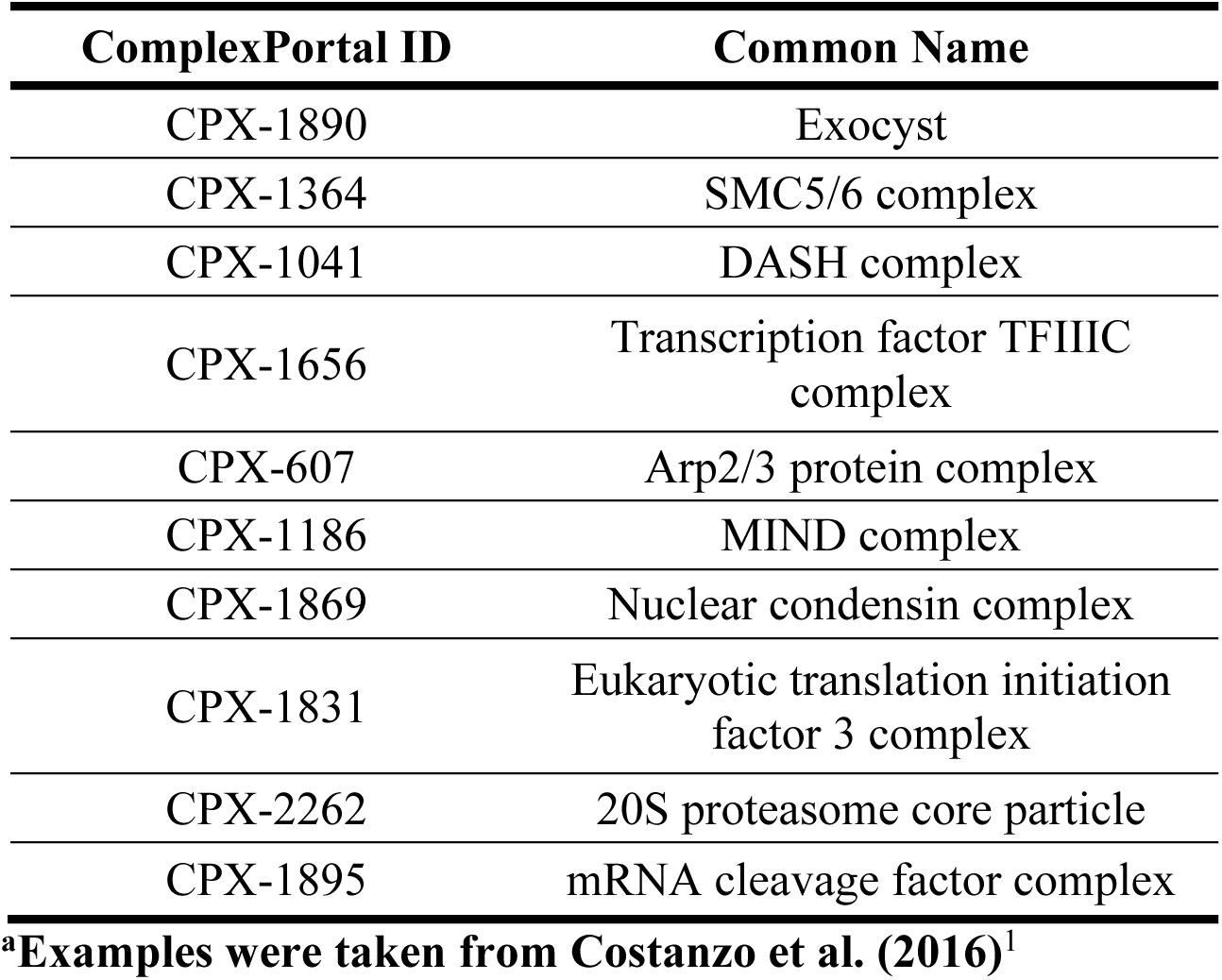
Essential Complexes Enriched for Negative GIs.

**Extended Data Table 2^a^:** Non-Essential Complexes Enriched for Positive GIs ComplexPortal ID Common Name.

| ComplexPortal ID | Common Name |
| --- | --- |
| CPX-779 | Elongator holoenzyme complex |
| CPX-1840 | Golgi transport complex |
| CPX-568 | CAF-1 complex |
| CPX-1197 | FAR complex |
| CPX-1342 | Set3C |
| CPX-1884 | HDA complex |
| CPX-1193 | Vacuolar proton-transporting V-type ATPase |
| CPX-1412 | Ubp3-Bre5 ubiquitin hydrolase complex |
| CPX-1655 | i-AAA complex |
| CPX-1178 | Cytoplasmic dynein complex |

<sup>a</sup>Examples were taken from Costanzo et al. (2016)<sup>1</sup>

**Extended Data Table 3:** Correlation coefficients between unbinned K_d_ and GI values.

| Organism | GI Type | Correlation Measure | Coefficient | p-value |
| --- | --- | --- | --- | --- |
| Yeast | Negative | Pearson | 0.22 | $1.3 \times 10^{-23}$ |
| | | Spearman | 0.24 | $1.1 \times 10^{-26}$ |
| | Positive | Pearson | 0.083 | $2.8 \times 10^{-3}$ |
| | | Spearman | 0.091 | $1.0 \times 10^{-3}$ |
| Human | Negative | Pearson | 0.098 | 0.11 |
|  |  | Spearman | 0.13 | 0.027 |
|  | Positive | Pearson | 0.038 | 0.48 |
|  |  | Spearman | 0.019 | 0.73 |

